# Transcriptome responses of two *Halophila stipulacea* seagrass populations from pristine and impacted habitats, to single and combined thermal and excess nutrient stressors, reveal local adaptive features and core stress-response genes

**DOI:** 10.1101/2025.01.21.634158

**Authors:** Hung Manh Nguyen, Beery Yaakov, Pedro Beca-Carretero, Gabriele Procaccini, Guannan Wang, Maheshi Dassanayake, Gidon Winters, Simon Barak

## Abstract

In their natural habitats, seagrasses face multiple abiotic stressors, which can often occur simultaneously. However, most studies investigating the effects of environmental stressors on seagrasses have focused on growth and physiological responses to single stressors. Here, we examined the transcriptome responses of the tropical seagrass *Halophila stipulacea* collected from a northern Gulf of Aqaba pristine site (South Beach - SB) and an anthropogenically-impacted site (Tur Yam - TY), grown in a mesocosm, and exposed to ecologically-relevant, single and combined, thermal and excess nutrient stressors. The combined thermal and nutrient stressor elicited greater transcriptome reprogramming than the single stressors in both populations and induced the expression of a combination-specific set of genes involved in abiotic and biotic stress responses. Furthermore, thermal stress exerted a more dominant influence than excess nutrient stress upon the transcriptome response to the combined stress. Transcriptomes of plants from the impacted TY site displayed reduced plasticity, the presence of genes exhibiting a “stress-ready” mode of expression under all stresses, and increased resilience (recovery to control transcriptomes). We also identified core stress-response genes that could be leveraged as early indicators of stress in the field. Overall, our data suggest that environmental conditions in seagrass habitats can drive local molecular adaptation, and that the response of seagrasses to combined stressors associated with climate change and coastal anthropogenic stressors cannot be predicted from the response to single stressors.

## Introduction

Seagrasses are marine flowering plants that thrive over thousands of kilometers of shallow-water shorelines from sub-Arctic to tropical regions (Short et al., 2007). Seagrasses support a wide range of vital ecosystem services, including oxygen production, provision of a habitat for numerous marine organisms such as fishes and invertebrates, water-quality improvement and nutrient recycling (Orth et al., 2006; Fourqurean et al., 2012; Nordlund et al., 2016; Lamb et al., 2017). They also represent one of the most significant blue-carbon sinks on Earth (Fourqurean et al., 2012; Macreadie and Hardy, 2018). In fact, seagrass meadows are ranked among the most valuable ecosystems with an estimated global value of US$ 2.8 million yr^-1^ km^-2^ (Costanza et al., 2014; Nordlund et al., 2016). Importantly, seagrass management and restoration have been recognized as a nature-based solution to tackle the impacts of global warming (Gattuso et al., 2018), supporting 16 out of 17 Sustainable Development Goals proposed by the United Nations (Unsworth et al., 2022).

Despite their crucial value, seagrasses are facing a global decline at an unprecedented rate of 110 km^2^ per annum due to both global stressors (e.g. ocean warming, marine heat waves, sea level rise) and local stressors (e.g. coastal development, eutrophication) (Orth et al., 2006; Waycott et al., 2009; Unsworth et al., 2014). Consequently, it has been estimated that 30% of the world’s seagrass meadows have been lost since 1980 (Waycott et al., 2009). This decline in seagrass meadows entails the loss of associated biodiversity, a decrease in primary productivity and local fishing grounds, and enhanced coastal erosion (Orth et al., 2006; Waycott et al., 2009; Unsworth et al., 2014). Taken together, the loss of these ecosystems will lead to severe ecological and socioeconomic consequences (Cullen-Unsworth et al., 2014; McKenzie et al., 2021; Turschwell et al., 2021).

The main concerns regarding the future of seagrass meadows in an era of ocean warming are increases in water temperatures (Nguyen et al., 2021) and the duration, severity, and frequency of anomalous heat waves (Brown, 2020). Indeed, in the past decade, many seagrass studies have turned their attention to studying the effects of thermal stress on different seagrass species, performing measurements at the molecular, physiological/biochemical, and even the ecosystem/planetary levels (Nguyen et al., 2021). However, it is clear that in their natural habitats, seagrasses encounter multiple stresses, which can often occur simultaneously. For example, eutrophication (Suonan et al., 2022) and the subsequent reduction in light availability may have a more substantial impact than changes in water temperature (Ralph et al., 2007; Premarathne et al., 2021). With ongoing, rapid increases in coastal human populations (Neumann et al., 2015), it is expected that in parallel to global warming, seagrass meadows will face continued exposure to stresses associated with coastal development. Indeed, it has been shown that in 72% of the sites where seagrass abundance has declined, the reduction was associated with indirect/direct coastal impacts that reduced water quality (Waycott et al., 2009).

There is a considerable body of research on the responses of terrestrial plants to a combination of stresses. These studies have revealed that responses to stress combinations are unique and cannot be predicted from plant responses to a single stress (Mittler and Blumwald, 2010; Zandalinas and Mittler, 2022). For example, *Arabidopsis thaliana* subjected to combined heat and drought exhibits high respiration, low photosynthesis, low stomatal conductance and high leaf temperature (Rizhsky et al., 2002; Rizhsky et al., 2004), whereas plants exposed to these stressors separately respond differently. In contrast, most stress-related seagrass studies have focused on the responses of seagrasses to a single, relatively short-term stress (Ralph, 1998; Shields et al., 2019; Nguyen et al., 2021). Nevertheless, a few studies have shown unique responses of seagrasses to stress combinations (Zimmerman et al., 2017; Mvungi and Pillay, 2019; Ontoria et al., 2019a; Ontoria et al., 2019b; Viana et al., 2020). For instance, *Posidonia oceanica* plants originating from pristine and eutrophic areas exhibit contrasting physiological and transcriptomic responses according to their location, and according to the application of single or combined stressors (Pazzaglia et al., 2020). Interestingly, a study that investigated the physiological responses of three co-existing seagrasses from Tanzanian coastal waters to single and combined thermal and nutrient stresses, showed that *Halophila stipulacea* is the most resilient species to all stress treatments (Viana et al., 2020).

The tropical seagrass *Halophila stipulacea* (Forsk.) Ascherson is a dominant component of the ecosystem in the northern Gulf of Aqaba [GoA], northern Red Sea. In this area, *H. stipulacea*, the most abundant and often the only seagrass species, forms monospecific beds within the soft sediments between the local coral reefs (Winters et al., 2020). Here, it has an outstanding wide depth distribution [2−50 m depth (Winters et al., 2017)] attributed to the clear waters in this region [light attenuation coefficient Kd _[400 to 700 nm]_ = 0.054 (Stambler, 2005)]. Due to its unique geographical location (surrounded by arid regions) and the bathymetric structure (i.e. The Straits of Tiran), the GoA is an oligotrophic ecosystem with a long water residence time of 3-8 years (Silverman and Gildor, 2008). It was recently reported that combined thermal and excess nutrient stressors caused a greater reduction in growth and higher mortality rates in two *H. stipulacea* northern GoA populations than either single stressor (Beca-Carretero et al., 2022). However, the effects of the combined stressor was dependent upon the seagrass population. Thus, in one population from a pristine site (denoted South Beach, SB), the thermal stressor alone caused an increase in growth. However, combined thermal and excess nutrient stressors severely reduced the positive effect of heat (i.e. excess nutrient stress [eutrophication] overrode thermal stress). In contrast, a second population (denoted Tur Yam, TY) situated in an anthropogenically-impacted site, exhibited almost no effect of the excess nutrient stressor upon plant growth whereas a combination of thermal and excess nutrient stressors had a deleterious effect on growth (i.e. thermal stress overrode nutrient stress). Furthermore, the impacted TY population exhibited lower and delayed mortality compared to plants from the pristine SB site when exposed to the combined stressor.

In the current study, we conducted a mesocosm experiment to investigate the transcriptome response of the two northern GoA *H. stipulacea* populations from the pristine and impacted sites, respectively (Beca-Carretero et al., 2022), to long-term, single and combined thermal and excess nutrient stressors. We show that the *H. stipulacea* transcriptome exhibits a unique response to the combined stressor and displays features that indicate molecular adaptations to site-specific environmental and anthropogenic stressors.

## Materials and Methods

### Sample collection areas

Plant samples were collected at two different monospecific meadows of *Halophila stipulacea* (South Beach and Tur Yam) in the northern Gulf of Aqaba, Eilat, Israel (**Fig. 1a,b**). The South Beach (SB; 29°29’34.2“N 34°54’21.6” E) site is located at the southern edge of the city of Eilat, far away from most hotels and urban areas, with clear water and high coral coverage. Therefore, the SB site is considered a pristine site (Beca-Carretero et al., 2022). Here, *H. stipulacea* meadows can be found at a depth range of 6 m to over 50 m along a steep bathymetric slope (∼18°) (Mejia et al. 2016). In contrast, the Tur Yam (TY; 29°30’58.0” N 34°55’41.6” E) site is 3 km away from the SB site and surrounded by hotels with medium-high anthropogenic pressures, a slightly turbid water column, and low coral coverage (Beca-Carretero et al., 2022). Thus, the TY site is considered a medium anthropogenically-impacted site (Mejia et al. 2016; Azcárate-García et al., 2020) and it has been shown that increased coastal development in Eilat has led to a rise in nutrient load in the water column (Nguyen et al., 2023). Here, *H. stipulacea* meadow depths range from 8 m to 12 m along a low bathymetric slope (∼5°) (Mejia et al., 2016; Rotini et al., 2017; Winters et al., 2017).

**Figure 1.**
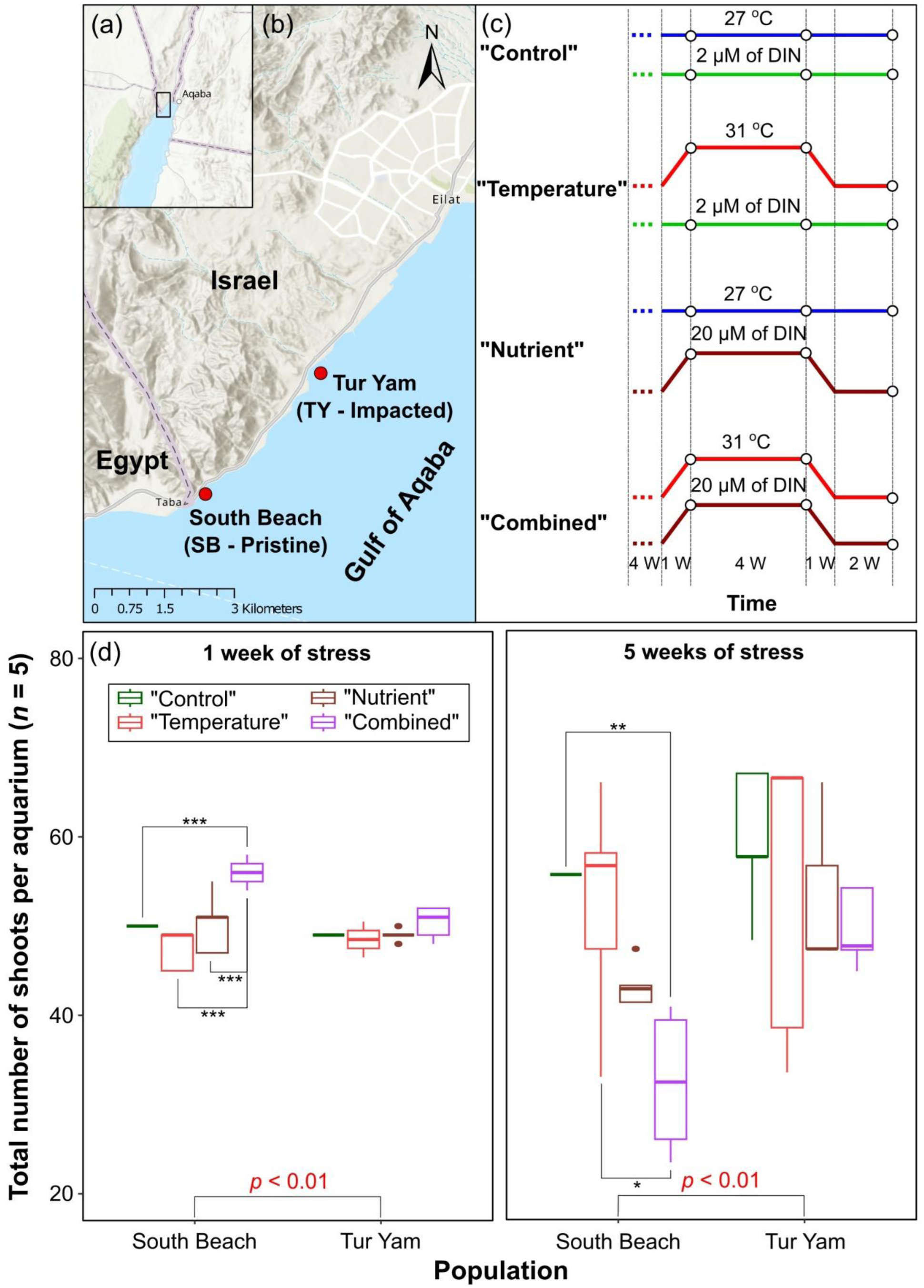
Sample collection sites, experimental scheme and the effect of single/combined thermal/excess nutrient stressors on the growth of SB (pristine site) and TY (impacted site) *H. stipulacea* plants. (a and b) Samples were collected at a pristine site and an anthropogenically-impacted site (red dots) along the Israeli coast in the Gulf of Aqaba; (c) Experimental scheme. Samples were collected for transcriptome analysis at three time points: one week of stress, five weeks of stress and two weeks of recovery. For details of experimental temperature and nutrient profiles see Materials and Methods, Supplementary Table S1 and Supplementary Fig. S1. Open circles, sampling time points for growth and RNA-seq; DIN, dissolved inorganic nitrogen; W, weeks; (d) TY shoot growth exhibits greater tolerance to single/combined thermal/excess nutrient stressors than SB plants. For box and whisker plots, the median (thick black line) and interquartile range (IQR) are shown. Whiskers indicate the maximum/minimum range. Closed circles demote extreme observations with values > 1.5 times the IQR. The influence of populations, treatments and their interactions on the total number of shoots at 1 week and 5 weeks of stress was evaluated using a two-way ANOVA, including the following factors: populations (North Beach vs. Tur Yam) and treatments (“Control” vs. “Temperature” vs. “Nutrient” vs. “Combined”). Before the analysis, the homogeneity of variance assumption was checked using Levene’s test while normality was assessed using the Shapiro-Wilk’s test. *p*-values at the bottom of each panel indicate significance difference between populations; **, *p* < 0.01 and ***, *p* < 0.001 (Tukey HSD *post hoc* test) indicate significance between control and treatment within a population.

### Sample collection

In June 2019, *H. stipulacea* plants were collected from both the SB and TY sites by SCUBA diving at a depth of 10 m. To reduce the likelihood of collecting the same clone twice, each *H. stipulacea* plant was collected 5-10 m away from each other. At both sites, irradiance level at the depth of sample collection was ∼250 μmol photons m^-2^ s^-1^ and water temperature was ∼26 ℃. These parameters were employed to mimic field conditions in the mesocosm system. Collected plants were kept in zip-lock plastic bags filled with natural seawater and quickly transported in cooler boxes to the seagrass mesocosm facility at the Dead Sea and Arava Science Centre (Hatseva, Israel) within less than 2 h post-collection.

### Experimental setup and design

Twenty aquaria (40 x 40 cm, 33 cm height) were filled with natural sediment collected from the shores of the GoA to a height of ∼7 cm and 40 L of artificial seawater (Red Sea Salt, Israel). *H. stipulacea* plants were gently cleaned of dead tissues and epiphytes. To standardize the experiment, 6-8 plants with a similar number of shoots (4-5 shoots) were selected from each of the SB and TY populations (i.e. 12 −16 plants in total) and planted in individual aquaria. Plants from the two populations were separated by a plastic divider, which allowed them to grow under the same conditions while avoiding any undesired population mixing.

Plants were allowed to acclimatize to the mesocosm conditions for 4 weeks before the start of the experiment. During this period, water temperature in all aquaria was kept at 27 ℃ which matches the annual sea surface temperature occurring in July when the experiment started (Nguyen et al., 2020). Irradiance level was set at ∼250 μmol photons m^-2^ s^-1^ with a 12 h:12 h light:dark cycle (similar to the irradiance in the field at the time of the experiment). Artificial seawater was generated using Red Sea Salt (Red Sea, Israel). The salinity level was maintained at 40 Practical Salinity Units (PSU). This salt concentration is similar to the salinity level of seawater at the collection sites and also yielded the control water nutrient concentration of ∼2 μM of Dissolved Inorganic Nitrogen (DIN).

For the duration of the experiment, salinity and water temperature were measured daily using the Multi 340i portable meter (WTW, Germany). Distilled water was added daily to compensate for water evaporation while 20% of the total water in the aquaria was renewed weekly to maintain the quality of the experimental seawater.

After acclimation, the 20 aquaria were assigned to four different treatments (*n* = 5 aquaria for each treatment) as depicted in **Fig. 1c**: (i) “Control” (27 ℃, 2 μM DIN); (ii) “Temperature” (31 ℃, 2 μM DIN); (iii) “Nutrient” (27 ℃, 20 μM DIN); (iv) “Combined” (31 ℃, 20 μM DIN). For the “Temperature” aquaria, water temperature was ramped up from 27 ℃ to 31 ℃ over a week (0.65 ℃ per day), kept at 31 ℃ for 4 weeks before being ramped down to 27 ℃ at the same rate of 0.65 ℃ per day, and then maintained at 27 ℃ for 2 weeks (**Supplementary Fig. S1**). Nutrient concentration in the “Temperature” aquaria was maintained at 2 μM DIN throughout the experiment. For the “Nutrient” aquaria, nutrient concentration was gradually increased to 20 μM DIN over a week by adding slow-release fertilizer pellets [Osmocote, 17:11:10 N:P:K, (Helber et al., 2021; Beca-Carretero et al., 2022)]. Ground pellets were dissolved in water and added three times per week on alternative days. Nutrient concentration was maintained for 4 weeks before being reduced to 2 μM DIN for a further 2 weeks. Water samples were collected weekly from each aquarium and nutrient levels were analyzed according to Beca-Carretero et al. (2022) (**Supplementary Table S1**). Water temperature of the “Nutrient” aquaria was kept at 27 ℃ throughout the experiment. For the “Combined” aquaria, both temperature and nutrient conditions were changed using the same schedule and method as for the “Temperature” and “Nutrient aquaria” (**Supplementary Fig. S1; Supplementary Table S1**). Salinity and light conditions of all aquaria throughout the experiment were maintained at the same level as during the acclimation period.

### RNA extraction and sequencing

*Halophila stipulacea* leaves from the 3^rd^ youngest shoot were collected at three different time-points: 1 week of stress, 5 weeks of stress and 2 weeks recovery (**Fig. 1c**). Harvested leaves were pooled from 6-8 plants from each aquarium. Epiphytes were carefully removed from the collected leaves and the cleaned material was flash-frozen in liquid nitrogen and stored at −80 ℃. Leaf samples were ground into a fine powder using a mortar and pestle, pre-cooled with liquid nitrogen. 50-100 mg of leaf power was used to extract total RNA with a Plant/Fungi Total RNA Purification Kit including on-column DNase I treatment (Norgen Biotek Corp., Canada) according to the manufacturer’s instructions. To prevent oxidation via phenolic compounds, 2% (w/v) polyvinylpyrrolidone-40 (PVP) was added to the lysis solution. RNA was sent to the Roy J. Carver Biotechnology Centre, University of Illinois (Urbana-Champaign, IL) for mRNA enriched polyA-selected library preparation and sequencing on an Illumina NovaSeq 6000 system to generate 150 nt paired-end reads.

### Transcriptome quantification and analysis

Raw sequencing reads (**Supplementary Table S2**) were first quality-verified using FastQC v0.11.8 (Andrews, 2010) and then mapped to the *H. stipulacea* genome (BioProject PRJNA1140917) using STAR v2.7.9a (Dobin and Gingeras, 2015). The average mapping rate across all samples was 74%. Differential expression was evaluated using DESeq2 v1.40.2 (Love et al., 2014) with a |log_2_ fold-change ≥ 1 and an adjusted *p*-value < 0.05. The mapped read counts were corrected using the ARSyNseq function in NOISeq v2.44.0 to minimize unknown batch effects and noise introduced from heterogeneous plants in the field (Tarazona et al., 2011). Functional annotation of protein coding transcripts was performed using Trinotate v4.0.0 (Bryant et al., 2017), with the following packages: TransDecoder v5.7.1, Diamond v2.1.9.163, Infernal 1.1.4, Signal v6.0g, tmhmm v2.0c, hmmer v2.3, and eggnog-mapper v2.1.10. A single representative transcript per gene in the reference genome was considered for differential expression analysis. The PCA plot was visualized on rlog-transformed counts. Differentially Expressed Genes (DEGs) were visualized using DiVenn 2.0 [available online at https://divenn.noble.org/index.php (Sun et al., 2019)]. Gene Ontology (GO) term enrichment of DEGs was performed using BiNGO v3.0.3 using a hypergeometric test with FDR correction and adjusted *p*-value < 0.05 (Benjamini and Hochberg, 1995) for over-represented GO terms in Cytoscape v3.10.0 (Otasek et al., 2019). Clustering of enriched GO terms was performed using GOMCL v0.0.1 (Wang et al., 2020a). Visualization of the clustered groups was performed in Cytoscape v3.10.0.

## Results

### Global analysis of *H. stipulacea* transcriptome responses to single and combined temperature and excess nutrient stresses

To examine the impact on the seagrass transcriptome of predicted stressors caused by global warming and excess nutrients from increased coastal development, we exposed the tropical seagrass *Halophila stipulacea* to single and combined temperature (heat) and excess nutrient stresses. Mesocosm-grown plants collected from pristine South Beach (SB) and anthropogenically -impacted Tur Yam (TY) sites (**Figs. 1a,b**) were challenged with an ecologically-relevant increase in water temperature (31 °C; “Temperature”) or an increase in nutrient concentration [20 μM dissolved inorganic nitrogen (DIN); “Nutrient”] or a “Combination” of the stresses (**Fig. 1c**). “Control” plants were maintained at 27 °C and 2 μM DIN. One week of “Combined” stress treatment led to a small increase in shoot growth (total number of shoots per aquarium) in the SB plants while TY plants were unaffected by any of the stressors (**Fig. 1d**). Five weeks of exposure to the “Temperature” stressor had no deleterious effect on SB plants but the “Nutrient” and “Combined” treatments led to a 23% and 42% reduction in shoot growth, respectively. In contrast, the single and combined stressors did not affect the shoot growth of TY plants. These data indicate that *H. stipulacea* plants from the anthropogenically-impacted TY site are more tolerant to stress than plants from the pristine SB site.

Analysis of the extent of global transcriptional adjustment in response to the single and combined stressors, showed that only the SB “Nutrient” transcriptome exhibited a significant increase in the median transcript abundance after one week of stress compared to the “Control” transcriptome (**Fig. 2a**). SB plants also displayed a significantly increased median transcript abundance after two weeks of recovery from the “Combined” treatment suggesting a possible delay in transcriptome recuperation. Similar results were observed when only genes associated with the GO-term “response to stress” were analyzed (**Supplementary Fig. S2**). Overall, these data indicate that transcriptomes from both SB and TY plants did not exhibit a large global transcriptional shift in response to stress. Therefore, we further investigated whether a smaller subset of transcripts was affected by the stressors using principle component analysis (PCA) of differentially expressed genes (DEGs). After 1 week of stress, transcriptomes of the “Temperature”, “Nutrient” and “Combined” samples from the pristine SB site were distinct from the “Control” samples (**Fig. 2b**). “Temperature” transcriptomes clustered near “Combined” samples suggesting that early during stress treatments, temperature stress exhibits a more dominant effect than nutrient stress when the plants are exposed to a combination of both stresses. In contrast, only the “Temperature” transcriptome was separated from “Control” samples in plants from the impacted TY site, while “Nutrient”, “Combined” and “Control samples clustered close to each other. After 5 weeks of stress, SB “Temperature” and “Combined” transcriptomes were positioned separately from the “Control” transcriptome with the “Combined’ transcriptome exhibiting the greatest difference. On the other hand, the “Nutrient” transcriptome clustered with the “Control” transcriptome. For plants from the TY site, transcriptomes from all stress treatments were distinct from the “Control” samples with the greatest separation observed for “Temperature” and “Combined” treatments that clustered close to each other. These data suggest a dominant role of temperature stress in the “Combined” treatment in plants from both sites after 5 weeks of stress.

**Figure 2.**
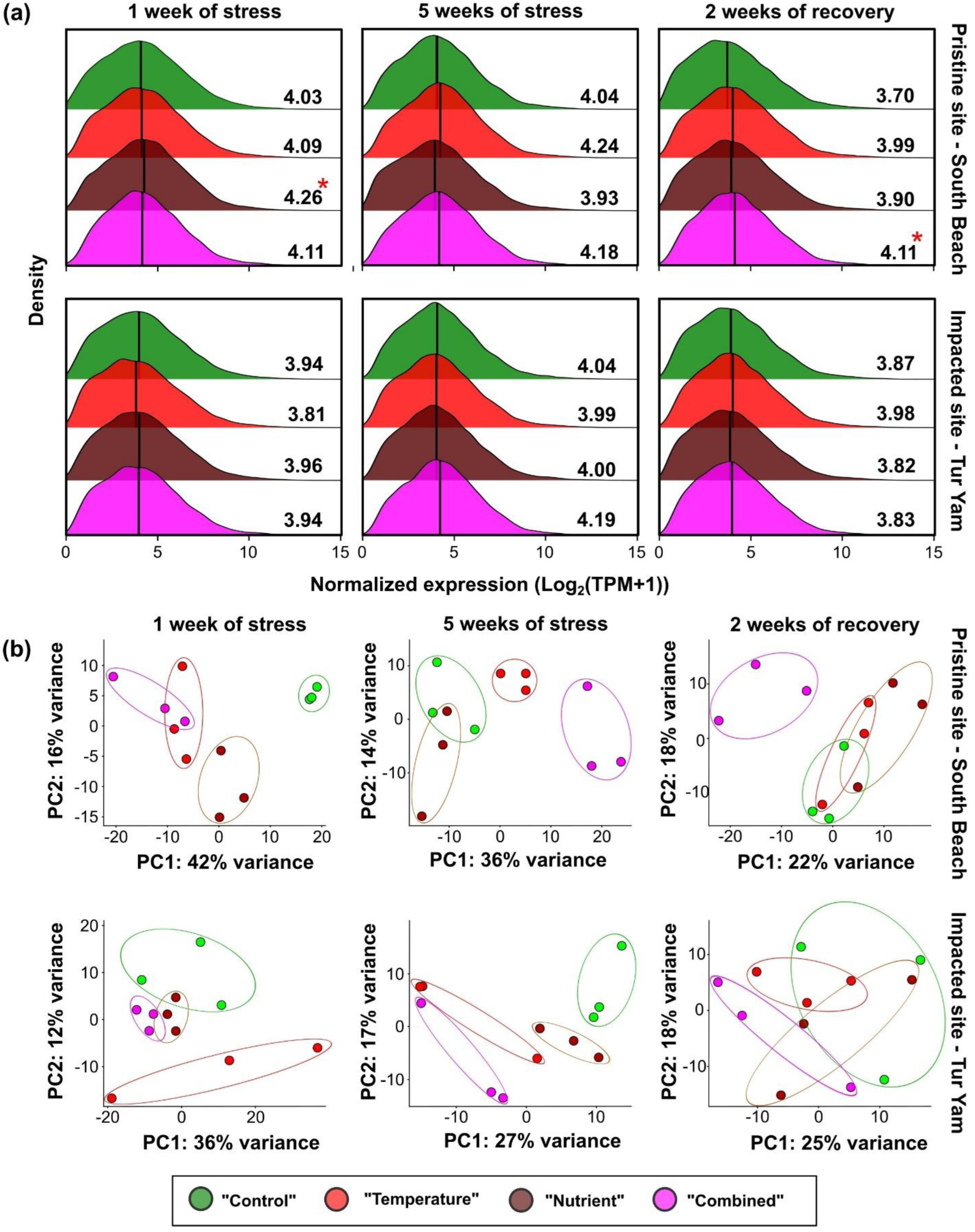
Global transcriptome response of SB (pristine site) and TY (impacted site) *H. stipulacea* plants to single and combined thermal and excess nutrient stresses. (a) Comparison of density distributions of transcript abundance between SB and TY plants subjected to control and stress treatments. Asterisks represent significant difference at *p* < 0.05 between the stress treatment compared to control; Normality was first tested using the Shapiro-Wilk test and homogeneity of variance was validated using the F-test. When assumptions for the Two Sample *t*-test were not met, the Wilcoxon rank-sum test was used. Black vertical line within plots is median expression; (b) Principal component analysis (PCA) of transcript profiles in control and stress-treated *H. stipulacea* SB and TY plants. PCA analysis was conducted on rlog-transformed read counts. Each point represents one biological replicate and the three replicates for each condition are depicted with the same colored symbol. Symbols are explained in the legend box and refer to the experimental scheme shown in Fig. 1c.

After 2 weeks of recovery from stress, the SB “Temperature” and “Nutrient” transcriptomes clustered with the “Control” samples suggesting that transcriptomes responding to the single stresses were recovering to their pre-stress states. In contrast, the “Combined” transcriptome was still distinct from the control samples even 2 weeks after the end of the stress treatment in agreement with the analysis of global transcriptome adjustment (**Fig. 2a**). These results indicate that a combination of temperature and nutrient stress has a more profound effect on the transcriptome than the single stresses. For TY plants, there was no clear separation of transcriptomes among any of the treatments, but it is not clear if this was due to the recovery of the stressed transcriptomes to a pre-stress state or large variation between replicates of each treatment. Overall, the density plots (**Fig. 2a**) and the PCA plots (**Fig. 2b**) indicate that while there was little overall shift in global transcriptome expression level in response to the stress treatments, a subset of genes did respond with a change in expression.

### TY plants display reduced transcriptome plasticity, possess “stress-ready” genes and exhibit greater transcriptome recovery after stress

To gain further insight into subsets of genes that exhibit different responses to the single and combined stressors between the pristine SB and impacted TY sites, we compared the numbers of SB and TY DEGs responding to each stress treatment (**Fig. 3a**; **Supplementary Tables S2 and S3**). Plants from the pristine SB site exhibited a greater number of total DEGs (i.e. the sum of DEGs for all the three treatments) than TY plants at all time-points. After one week of stress, there were 3.7-fold more DEGs in the SB plants (1,744 SB vs. 465 TY) while after 5 weeks of stress, there were 1.7-fold more DEGs in SB plants (1,577 SB vs. 918 TY). The larger number of SB DEGs in response to stress was also reflected after the 2-week recovery period where 3.7-fold more DEGs were observed in SB plants compared to TY plants (1,042 SB vs. 281 TY). A similar trend of more SB DEGs than TY DEGS was also observed for each individual stress treatment. This was particularly striking after one-week’s excess nutrient stress where there were 21.5-fold more DEGs in plants from the pristine SB site than from the impacted TY site (560 SB vs. 26 TY).

These data suggest a difference in plasticity of gene expression between the two populations. We therefore analyzed transcriptome plasticity in terms of reaction norms i.e. the pattern of transcriptome expression (DEGs) of each population across the various stress treatments (Kusmec et al., 2018; Rivera et al., 2021). This analysis revealed three main differences between the two populations (**Fig. 3b**): (i) for every stressor and at every time point, SB plants exhibited a greater number of DEGs inferring greater transcriptome plasticity than in TY plants; (ii) each population exhibited a different pattern of transcriptome plasticity in response to each stressor suggesting a genotype x environment interaction. For instance, TY transcriptomes exhibited a peak number of DEGs after 5 weeks of stress in response to all stress treatments whereas the SB transcriptomes exhibited stress-specific patterns of DEGs; (iii) TY stress-induced transcriptomes exhibited greater recovery to near “Control” transcriptomes across all stress treatments compared to SB transcriptomes.

**Figure 3.**
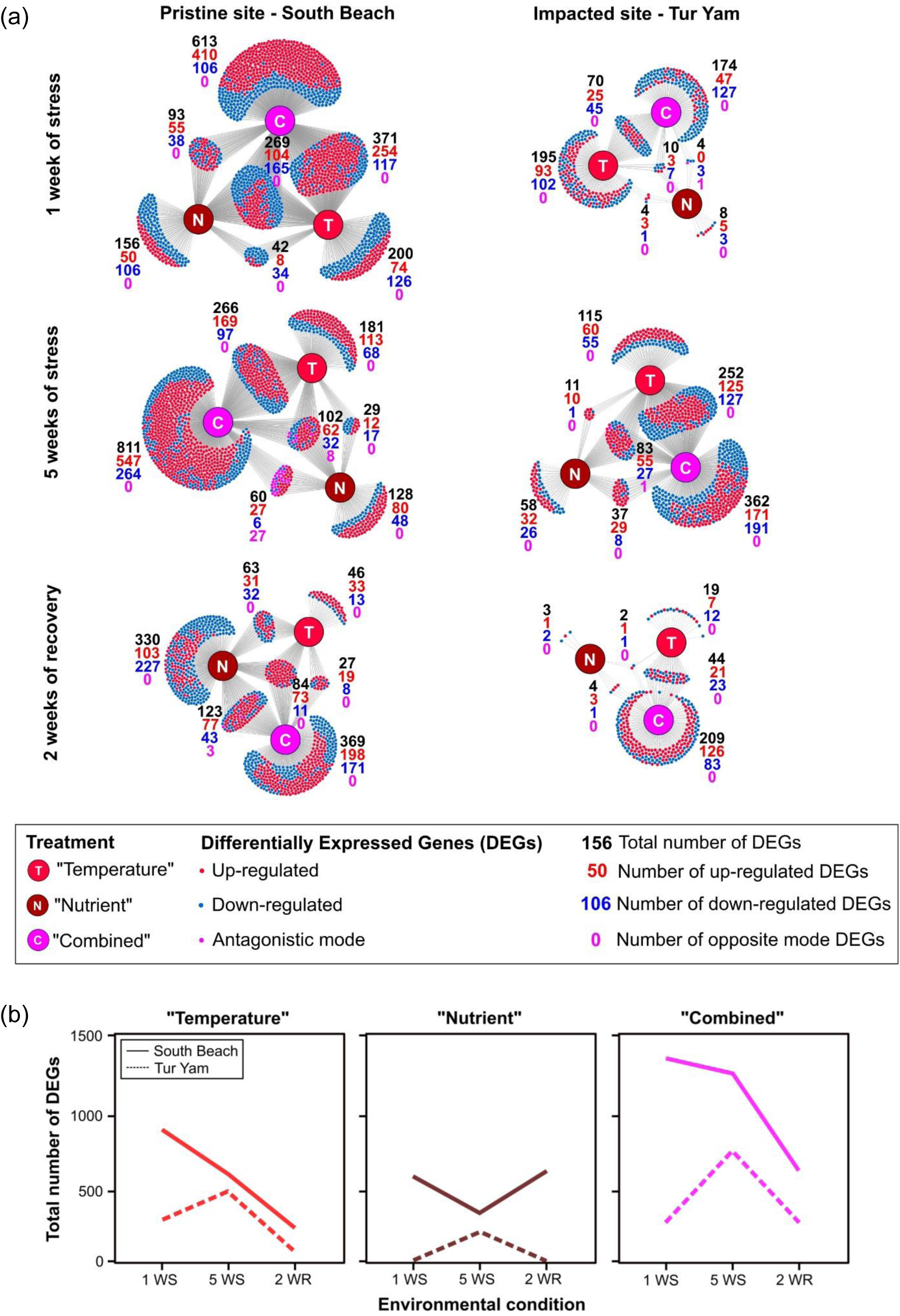
Comparison of differentially expressed genes (DEGs) and transcriptome plasticity between SB (pristine site) and TY (impacted site) *H. stipulacea* plants in response to single and combined thermal and excess nutrient stresses. (a) TY plants display a more restrained stress-mediated transcriptome response and both SB and TY plants exhibit stress combination-unique DEGs. Divenn diagrams show common and unique DEGs across the three treatments for each population at each time point. Each dot represents one DEG and is coloured according to its DEG mode (i.e. up-regulated, down-regulated or opposite mode) (Supplementary Tables S2 and S3); (b) Transcriptome plasticity of SB and TY *H. stipulacea* plants in response to single/combined thermal/excess nutrient stresses and during recovery. Plasticity was assessed as reaction norms of transcriptome expression (no. of DEGs) over all time points for each stress treatment.

The reduced plasticity of the TY transcriptome is reminiscent of Brassicaceae halophytic models whose transcriptomes exist in a “stress-ready” state. Here, genes whose expression is up/downregulated in response to ionic stress in stress-sensitive *Arabidopsis thaliana* display high or low constitutive expression in the halophytes even under stress-neutral conditions (Kazachkova et al., 2018; Wang et al., 2021). We therefore examined whether TY plants displayed signatures of a “stress-ready” transcriptome. DEGs from both populations were assigned to two idealized transcriptional response modes: “Stress-ready” or “Shared response” (where expression of genes is either up/downregulated in both SB and TY plants) (**Fig. 4; Supplementary Table S4**). Of the TY genes assigned to these response modes, 47%, 92% and 46% exhibited a “stress-ready” mode in the “Temperature”, “Nutrient” and “Combined” treatments, respectively. In contrast, no “stress-ready” response modes were observed in the SB “Temperature”, or “Nutrient” samples, while only 0.7% (2 genes) displayed a “stress-ready” mode in the “Combined” samples.

**Figure 4.**
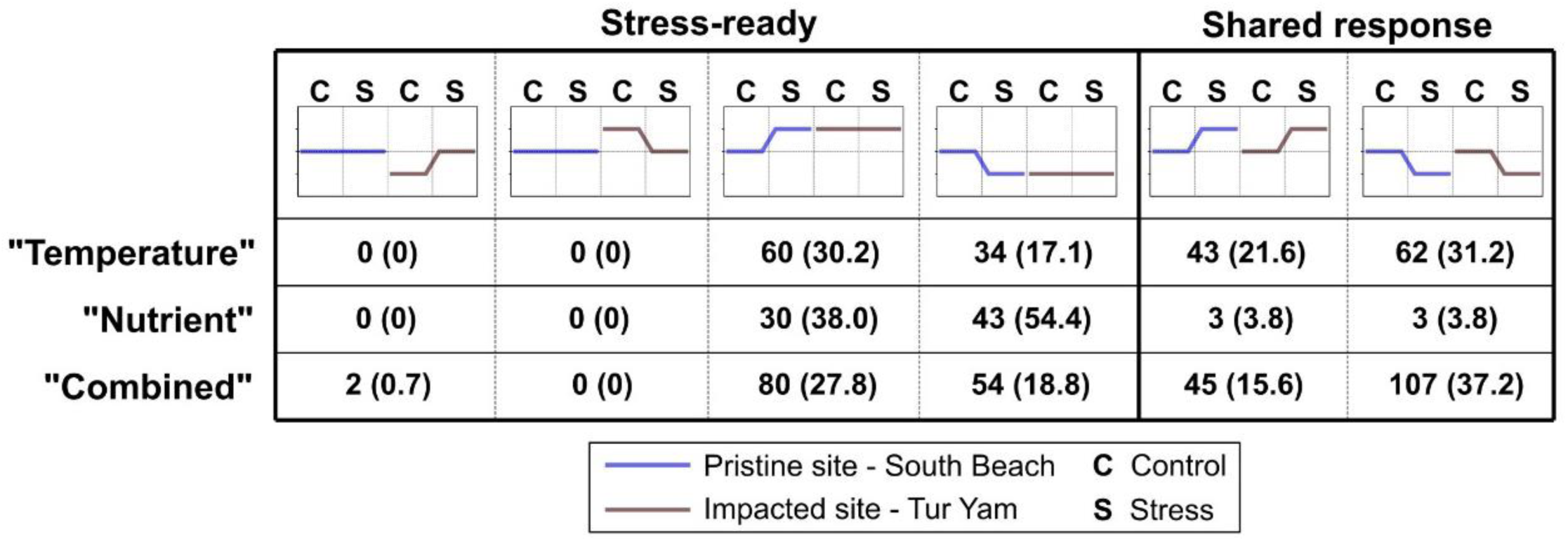
The TY transcriptome possesses genes exhibiting a “stress-ready” mode of expression. The DiVenn diagrams (Fig. 3a) were used to identify DEGs that were: (i) up-/down-regulated in SB plants but were constitutively up-/down-regulated in TY and vise-versa (“Stress-ready”) or (ii) exhibited up-/down regulation in both populations (“Shared response”) under the various stress treatments. Differences in absolute transcripts levels between control samples were identified by comparing transcripts per kilobase million (TPM) expression values (Student’s *t*-test, *p* < 0.05).

Constitutively up-regulated “stress-ready” TY genes from the “Temperature” and “Combined” samples were both enriched in biological processes related to oxidative stress and cell wall remodeling, while constitutively down-regulated “stress-ready” genes were enriched in GO-terms associated with regulation of transcription (**Table 1; Supplementary Table S5**). “Stress-ready” up-regulated TY genes from the “Nutrient” samples were only enriched in ethylene-mediated signaling (2 genes) and down-regulated “stress-ready” genes were not enriched in any GO-terms. While only TY transcriptomes exhibited “stress-ready” genes, we observed that 53% of genes showed a “Shared Response” mode between the two species in the “Temperature” and “Combined” samples and 3.8% (3 genes) exhibited this mode of expression in the “Nutrient” samples. Taken together, our findings showing a relatively restrained response of the TY transcriptome to stress and the presence of “stress-ready” genes only in TY plants, support the notion that TY plants from the impacted site exist in a “stress-ready” state.

**Table 1:**
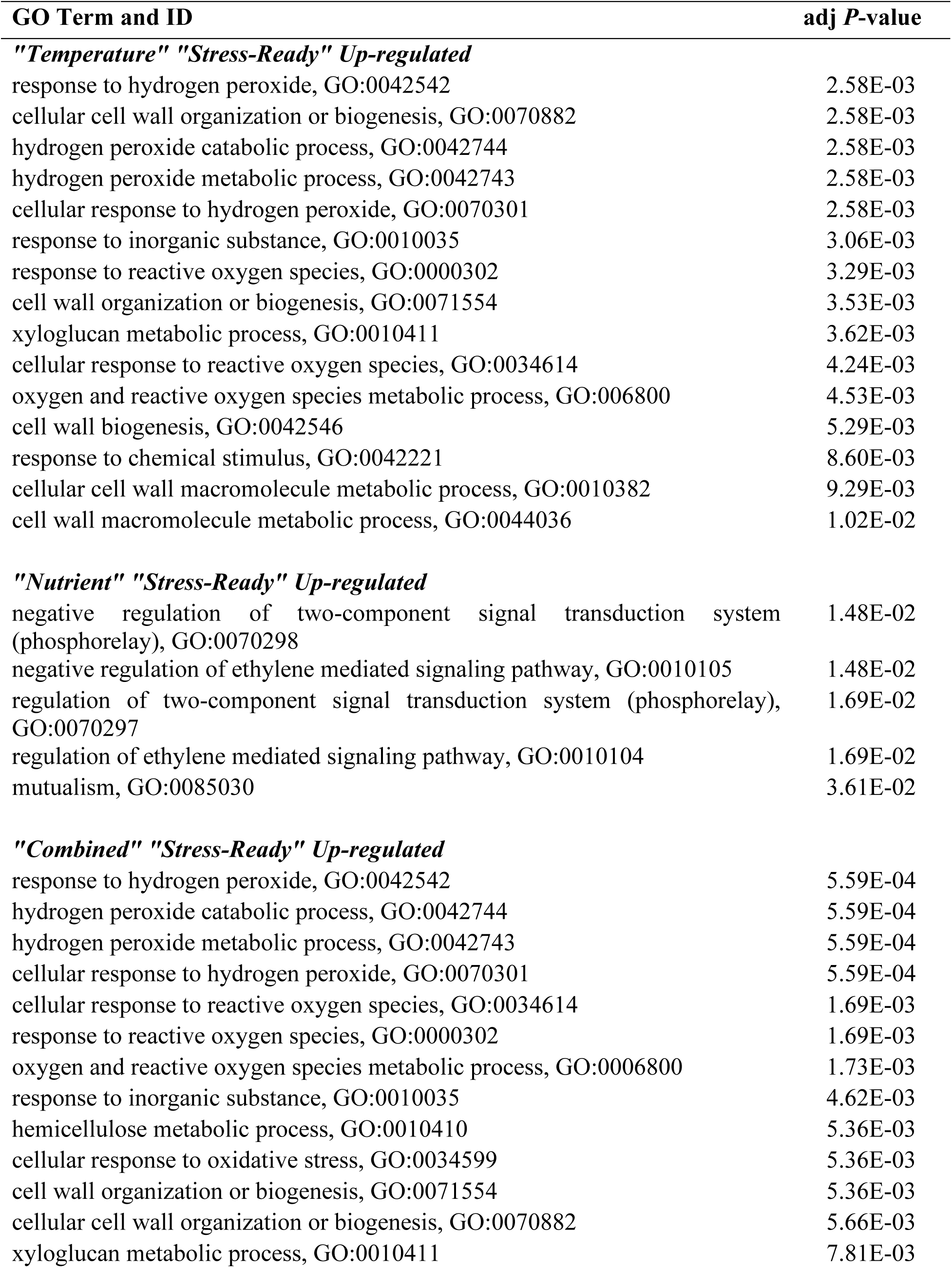
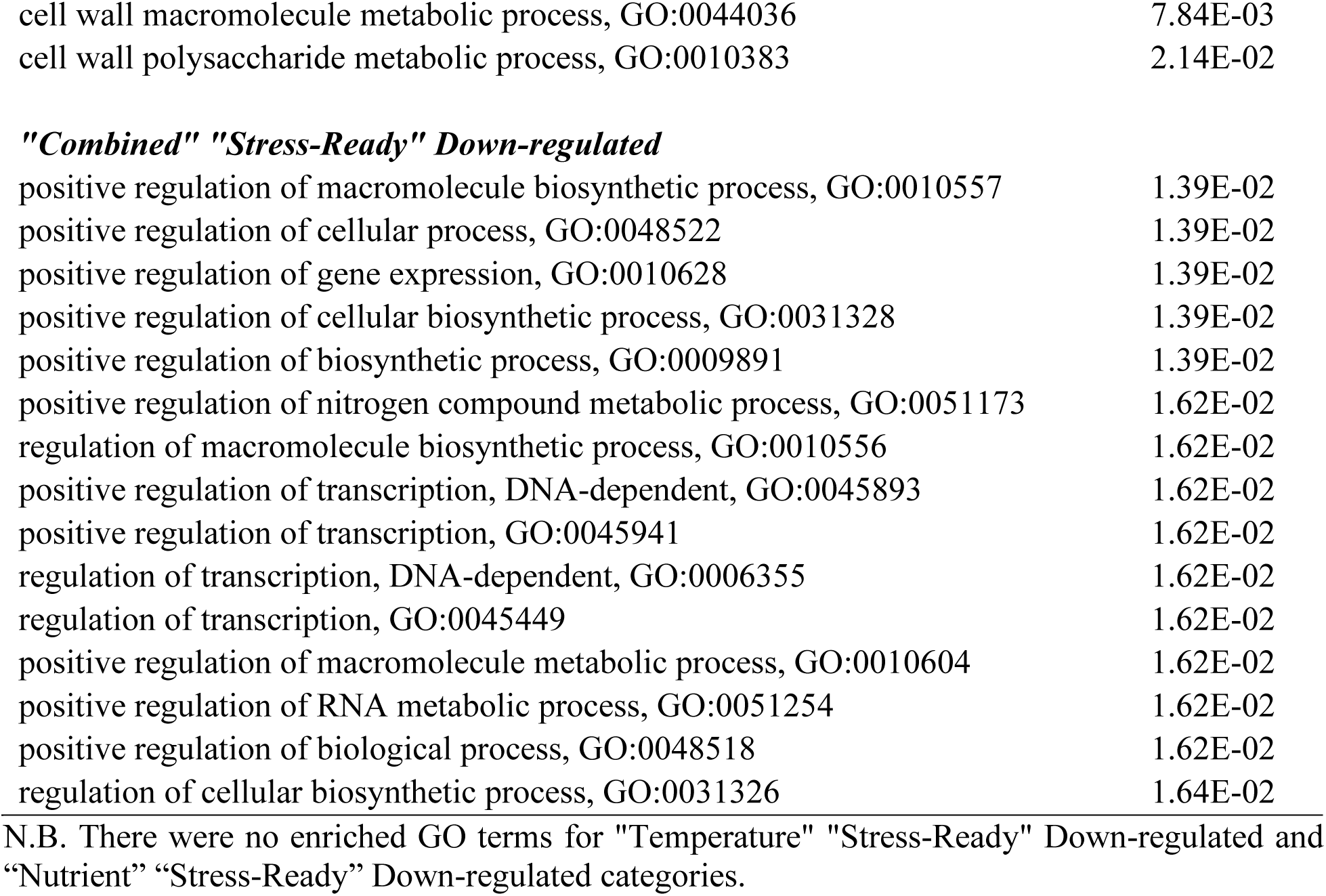
Top 15 biological process GO-terms for “stress-ready” TY genes.

| GO Term and ID | adj *P*-value |
| --- | --- |
| ***"Temperature" "Stress-Ready" Up-regulated*** |  |
| response to hydrogen peroxide, GO:0042542 | 2.58E-03 |
| cellular cell wall organization or biogenesis, GO:0070882 | 2.58E-03 |
| hydrogen peroxide catabolic process, GO:0042744 | 2.58E-03 |
| hydrogen peroxide metabolic process, GO:0042743 | 2.58E-03 |
| cellular response to hydrogen peroxide, GO:0070301 | 2.58E-03 |
| response to inorganic substance, GO:0010035 | 3.06E-03 |
| response to reactive oxygen species, GO:0000302 | 3.29E-03 |
| cell wall organization or biogenesis, GO:0071554 | 3.53E-03 |
| xyloglucan metabolic process, GO:0010411 | 3.62E-03 |
| cellular response to reactive oxygen species, GO:0034614 | 4.24E-03 |
| oxygen and reactive oxygen species metabolic process, GO:006800 | 4.53E-03 |
| cell wall biogenesis, GO:0042546 | 5.29E-03 |
| response to chemical stimulus, GO:0042221 | 8.60E-03 |
| cellular cell wall macromolecule metabolic process, GO:0010382 | 9.29E-03 |
| cell wall macromolecule metabolic process, GO:0044036 | 1.02E-02 |
| ***"Nutrient" "Stress-Ready" Up-regulated*** |  |
| negative regulation of two-component signal transduction system (phosphorelay), GO:0070298 | 1.48E-02 |
| negative regulation of ethylene mediated signaling pathway, GO:0010105 | 1.48E-02 |
| regulation of two-component signal transduction system (phosphorelay), GO:0070297 | 1.69E-02 |
| regulation of ethylene mediated signaling pathway, GO:0010104 | 1.69E-02 |
| mutualism, GO:0085030 | 3.61E-02 |
| ***"Combined" "Stress-Ready" Up-regulated*** |  |
| response to hydrogen peroxide, GO:0042542 | 5.59E-04 |
| hydrogen peroxide catabolic process, GO:0042744 | 5.59E-04 |
| hydrogen peroxide metabolic process, GO:0042743 | 5.59E-04 |
| cellular response to hydrogen peroxide, GO:0070301 | 5.59E-04 |
| cellular response to reactive oxygen species, GO:0034614 | 1.69E-03 |
| response to reactive oxygen species, GO:0000302 | 1.69E-03 |
| oxygen and reactive oxygen species metabolic process, GO:0006800 | 1.73E-03 |
| response to inorganic substance, GO:0010035 | 4.62E-03 |
| hemicellulose metabolic process, GO:0010410 | 5.36E-03 |
| cellular response to oxidative stress, GO:0034599 | 5.36E-03 |
| cell wall organization or biogenesis, GO:0071554 | 5.36E-03 |
| cellular cell wall organization or biogenesis, GO:0070882 | 5.66E-03 |
| xyloglucan metabolic process, GO:0010411 | 7.81E-03 |
| cell wall macromolecule metabolic process, GO:0044036 | 7.84E-03 |
| cell wall polysaccharide metabolic process, GO:0010383 | 2.14E-02 |
| ***"Combined" "Stress-Ready" Down-regulated*** |  |
| positive regulation of macromolecule biosynthetic process, GO:0010557 | 1.39E-02 |
| positive regulation of cellular process, GO:0048522 | 1.39E-02 |
| positive regulation of gene expression, GO:0010628 | 1.39E-02 |
| positive regulation of cellular biosynthetic process, GO:0031328 | 1.39E-02 |
| positive regulation of biosynthetic process, GO:0009891 | 1.39E-02 |
| positive regulation of nitrogen compound metabolic process, GO:0051173 | 1.62E-02 |
| regulation of macromolecule biosynthetic process, GO:0010556 | 1.62E-02 |
| positive regulation of transcription, DNA-dependent, GO:0045893 | 1.62E-02 |
| positive regulation of transcription, GO:0045941 | 1.62E-02 |
| regulation of transcription, DNA-dependent, GO:0006355 | 1.62E-02 |
| regulation of transcription, GO:0045449 | 1.62E-02 |
| positive regulation of macromolecule metabolic process, GO:0010604 | 1.62E-02 |
| positive regulation of RNA metabolic process, GO:0051254 | 1.62E-02 |
| positive regulation of biological process, GO:0048518 | 1.62E-02 |
| regulation of cellular biosynthetic process, GO:0031326 | 1.64E-02 |

N.B. There were no enriched GO terms for "Temperature" "Stress-Ready" Down-regulated and "Nutrient" "Stress-Ready" Down-regulated categories.

### The “Combined” stress treatment induces a higher number of DEGs and expression of a unique combination-specific set of genes

Our data show that the “Combined” treatment led to a greater overall number of DEGs than either of the single stresses in both populations and at each time point (**Fig. 3a; Supplementary Tables S2 and S3**). For example, after 1 week of stress, SB plants exhibited 560, 882 and 1,346 DEGs for “Nutrient”, Temperature” and “Combined”, respectively. Importantly, we found a large proportion of stress combination-specific DEGs. Thus, at one week of stress, 45.5% and 67.4% of the total number of DEGs were unique to the “Combined” treatment for the SB and TY plants, respectively. At five weeks of stress, 65.4% and 49.3% of SB and TY DEGs, respectively, were combination-specific while 61.2% and 80.7% of SB and TY DEGs, respectively, were unique to plants exposed to the “Combined” treatment after two weeks of recovery. We also detected DEGs that were common to different stress treatments but that exhibited an antagonistic mode of expression (i.e. up-regulated in the single stress but down-regulated in the combined stress or vice-versa). For instance, we found 27 DEGs common to the “Nutrient” and “Combined” treatments, and 8 DEGs common to all three treatments that displayed an antagonistic mode of expression in SB plants after 5 weeks of stress.

To examine the functional significance of the combination-specific DEGs, we performed GO-term overrepresentation analyses on these gene sets (**Fig. 5**; **Supplementary Tables S6-S9**). The results revealed that for both populations after 1-week and 5 weeks of stress, combination-specific DEGs were enriched in biological processes related to “response to abiotic stimulus” including water, salt, oxidative, hypoxia and cold stresses, and the response to abscisic acid. “Defense response” including response to fungi, bacteria and insects (response to wounding, chitin, jasmonic acid, oxylipin metabolic process), and “developmental process” (development of shoots and reproductive structures) were also enriched in both populations and at both time points. Signatures of the response to excess nutrient stress (increased DIN) were manifested in GO-terms such as “nitrate transport” and “ion transport” (including nitrate transporters” and “amino acid transport”). In addition, the gene encoding HY5, a master transcriptional regulator controlling processes such as photomorphogenesis and nitrate acquisition (Xiao et al., 2022), was observed in both populations under enriched GO-terms “regulation of macromolecule metabolic process”, “regulation of developmental process”.

**Figure 5.**
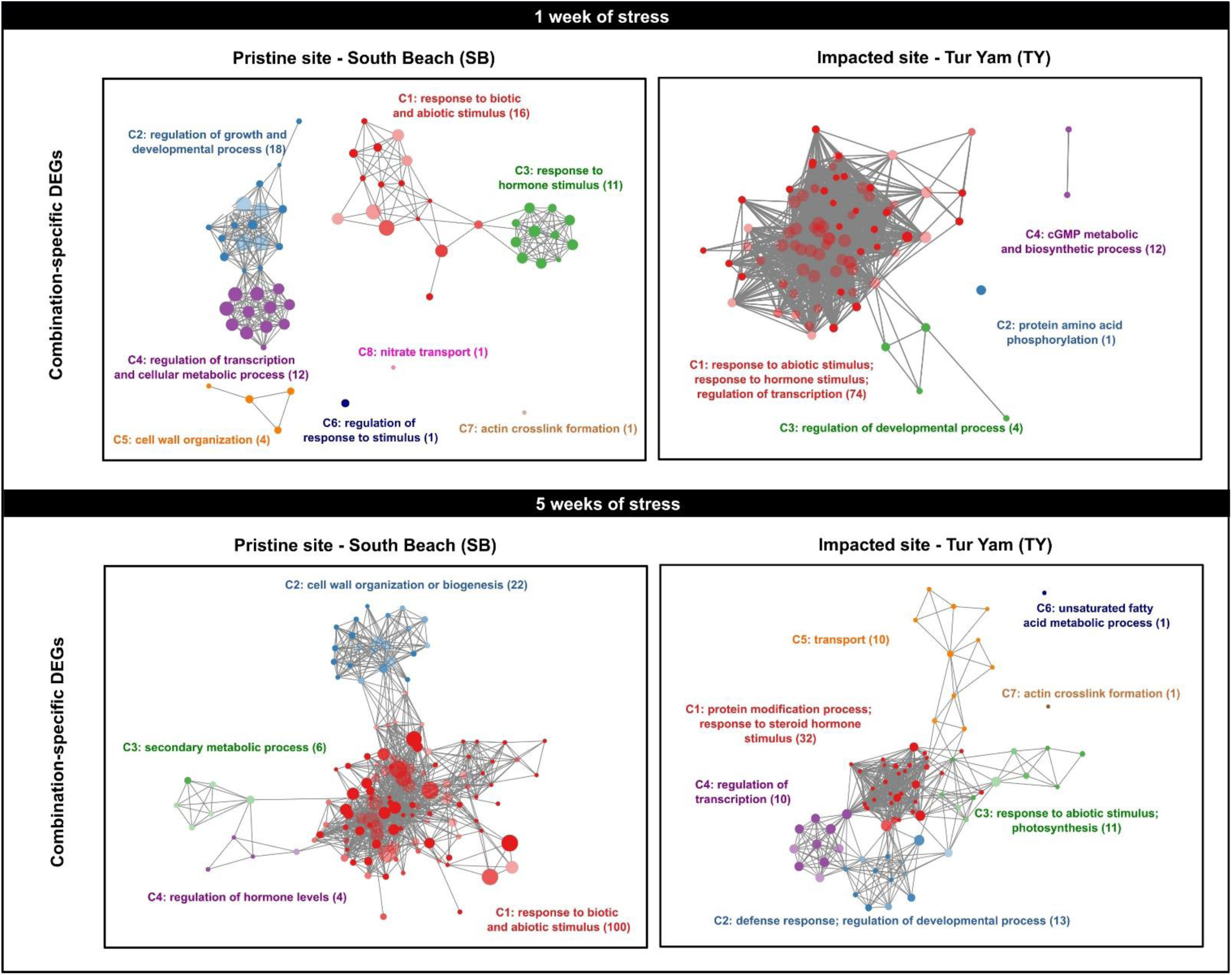
Functional clusters enriched in combined stress-specific DEGs at one and five weeks of stress for SB (pristine site) and TY (impacted site) *H. stipulacea* plants. For full list of GO-terms in each cluster and gene lists for selected GO-terms, see Supplementary Tables S6 to S9. Clustering was performed with the GOMCL tool (https://github.com/Guannan-Wang/GOMCL) (Wang et al., 2020a) Clusters are colored differently and labeled with the representative functional term. Each node represents a GO term and node size signifies the number of genes in the test set assigned to that functional term; the number of genes in each cluster is in parentheses. The shade of each node represents the *p*-value assigned by the enrichment test (false discovery rate (FDR)-adjusted *p* < 0.05) with darker shades indicating smaller *p*-values. GO-terms sharing > 50% of genes are connected by edges. Only selected clusters are highlighted, the rest are grayed out.

### Identification of Core Stress-Response genes

Our data presented the opportunity to identify genes that are part of the core stress machinery (Van den Broeck et al., 2017); i.e. they are differentially expressed in response to stress, irrespective of population and type of stress. Such genes could be used as molecular markers of stress in *H. stipulacea* plants before growth and physiological phenotypes are visible. We therefore pinpointed DEGs that were common to plants from the SB and TY sites in response to each stress treatment (**Fig. 6a**). Genes common to those three gene sets were then selected as putative *Core Stress-Response* (*CSR*) genes. We identified 38 *CSR* genes (**Fig. 6b**) including genes involved in nutrient (nitrogen, phosphate, potassium) responses (Liu et al., 2009; Huang et al., 2016; Sato et al., 2017; Ueda et al., 2020; Liu et al., 2023a), stress responses (Blanvillain et al., 2009; Krishnaswamy et al., 2011; Luhua et al., 2013; Fu et al., 2014; Liu et al., 2016; He et al., 2021; Huo et al., 2021; Zhang et al., 2023; Méndez-Gómez et al., 2024), the circadian clock (Rawat et al., 2009; Dai et al., 2011) and defense (Bhattarai et al., 2010; Wang et al., 2020b; Huo et al., 2021) among other biological functions related to developmental processes (Kubo et al., 1999; Woo et al., 2007; Martín et al., 2013; Yang et al., 2022), germination (Bassel et al., 2011; Li et al., 2012), transcriptional regulators (Reyes et al., 2004), and root-specific functions (Forsthoefel et al., 2005).

**Figure 6.**
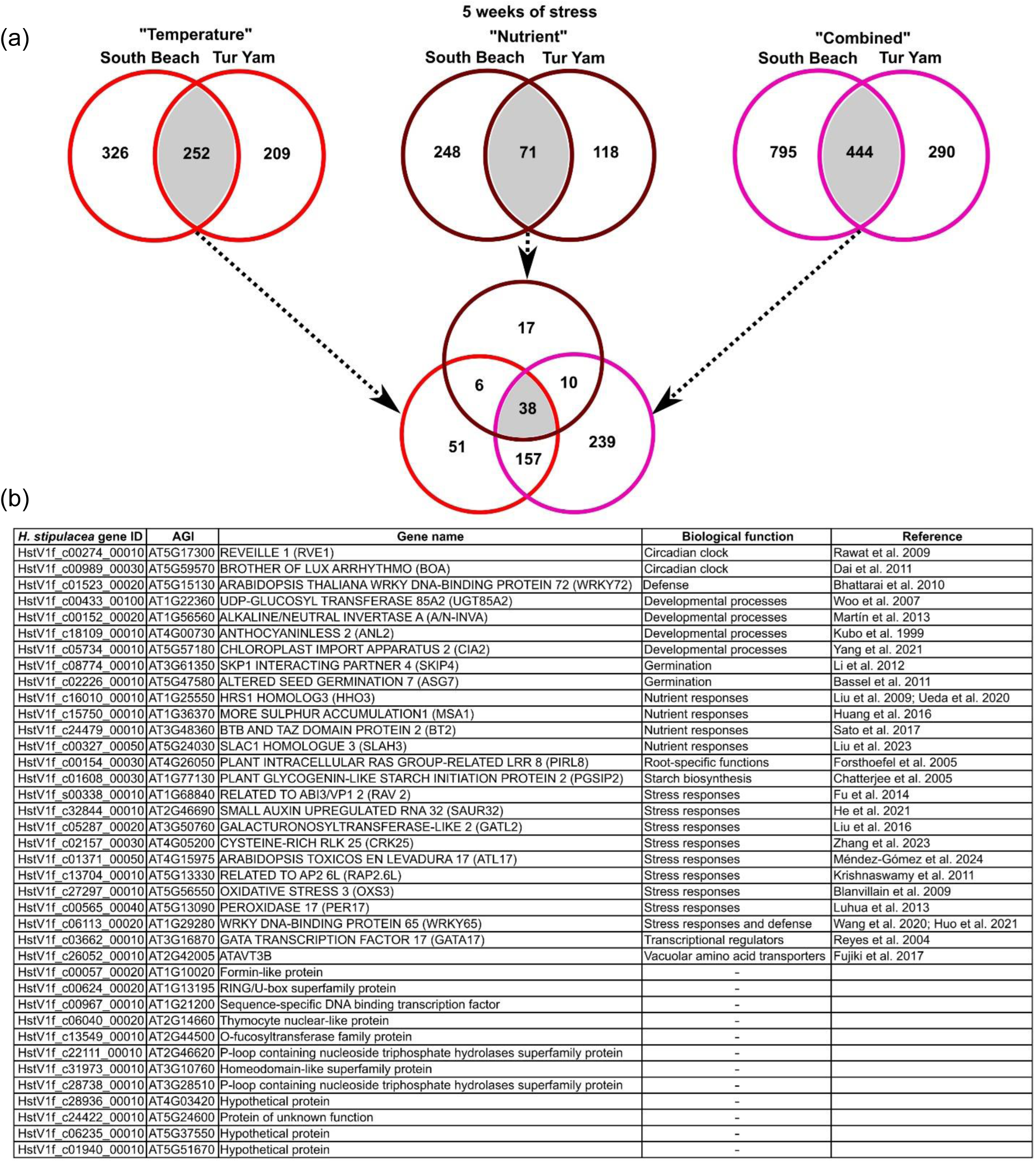
Core Stress-Response (*CSR*) genes. (a) Scheme for identifying *CSR* genes; (b) List of *CSR* genes and their biological functions.

## Discussion

In this study, we examined the behavior of the *Halophila stipulacea* transcriptome when challenged by single and combined heat and excess nutrient stressors and investigated whether this response was dependent upon site-specific acclimation to local conditions. The SB and TY *H. stipulacea* populations are located in pristine and anthropogenically-impacted sites (**Fig.1a,b**), respectively, and we showed that after 5 weeks of stress, TY plants exhibited no reduction in shoot growth in response to any of the stress treatments (**Fig. 1d**). On the other hand, the “Nutrient” and “Combined” treatments led to decreased shoot growth in SB plants. Taken together with data showing that the TY plants display lower and delayed mortality in response to combined thermal and excess nutrient stressors compared to SB plants (Beca-Carretero et al., 2022), our results suggest that plants from the TY population are more tolerant to stress.

### Thermal stress has a greater role than nutrient stress in governing the “Combined” transcriptome

PCA indicated that thermal stress plays a more dominant role than excess nutrient stress in shaping the *H. stipulacea* transcriptome under combined stress conditions (**Fig. 2b**). This result is further supported by findings that the “Temperature” transcriptomes possess many more DEGs in common with the “Combined” transcriptomes than between the “Nutrient” and “Combined” transcriptomes **(Fig. 3a**). Thermal stress also induces the expression of many more genes than an increase in nutrient load (**Fig. 3a**). At the level of growth response, Viana et al. (2020) showed that temperature is a stronger stressor than nutrient stress in *H. stipulacea* from an East African population although we did not observe such an effect on SB or TY plants growth in this study (**Fig. 1d**). A similar dominance of thermal stress over high nutrient stress on growth and physiological parameters was observed in the seagrass *Cymodocea nodosa* (Ontoria et al., 2019b). On the other hand, nutrient increase has a stronger effect than thermal stress on the performance of the temperate seagrass species *Zostera capensis* and *Posidonia oceanica* (Mvungi and Pillay, 2019; Pazzaglia et al., 2020). In the case of *P. oceanica*, the stronger effect of high nutrient load is also borne out at the transcriptome level (Pazzaglia et al., 2022). Overall, these data indicate species-specific and region/habitat-specific differences in the response to various stressors.

### The “Combined” stress treatment elicits greater transcriptome reprogramming than the single stresses and induces combination-specific expression of genes involved in stress responses

The combination of thermal and excess nutrient stressors clearly had a greater effect on transcriptome reprogramming than either of the single stresses. The “Combined” transcriptome exhibited, in general, the greatest separation from the “Control” transcriptome in the PCA plots. Moreover, at least in the SB population, the “Combined” transcriptome did not fully recover to its pre-stress state upon removal of the stressors (**Fig. 2b**). Furthermore, the “Combined” treatment induced a greater number of DEGs than either of the single stressors (**Fig. 3a**). These data are consistent with a greater reduction in *H. stipulacea* growth and increased mortality caused by the combined thermal and excess nutrient stressors compared to the single stressors (Beca-Carretero et al., 2022). Such a synergistic effect of combined stressors leading to decreased plant growth and enhanced transcriptome response has also been observed in terrestrial plants (Suzuki et al., 2016; Shaar-Moshe et al., 2017; Balfagón et al., 2019). However, some stress combinations elicit an antagonistic effect, which can be species-dependent (Mittler and Blumwald, 2010; Zandalinas and Mittler, 2022). In the case of a combination of thermal stress and increased nutrient loads, a rise in nitrogen application in plant species such as tomato and rice reduces the deleterious effects of heat stress (Luo et al., 2023; You et al., 2023). Conversely, and similar to *H. stipulacea* in the current study, increased nitrogen application exacerbates the adverse effects of heat stress in maize (Ordóñez et al., 2015).

Our data also revealed a large proportion of “Combination”-unique DEGs (**Fig. 3a; Supplementary Tables S2 and S3**). Similarly, studies with various terrestrial plants and, more recently, with the seagrass *P. oceanica* have also discovered unique stress combination-specific DEGs (Rizhsky et al., 2002; Rizhsky et al., 2004; Atkinson et al., 2013; Rasmussen et al., 2013; Sewelam et al., 2014; Suzuki et al., 2016; Shaar-Moshe et al., 2017; Balfagón et al., 2019; Zandalinas et al., 2020; Zandalinas et al., 2021; Pazzaglia et al., 2022; Tan et al., 2023). The complexity of plant responses to stress combinations is also exemplified by the finding that both common genes shared by several stresses and combination-unique genes, exhibit different gene expression patterns (e.g. antagonistic, additive) depending upon the stress combination (Rasmussen et al., 2013; Sewelam et al., 2014; Shaar-Moshe et al., 2017; Zandalinas and Mittler, 2022). Thus, the transcriptome response of plants, including temperate and tropical seagrasses, to combinations of stressors, cannot be predicted from the response to single stressors. In keeping with terrestrial plants, “Combination”-specific DEGs in *H. stipulacea* were enriched in biological processes related to the response to various abiotic stresses (**Fig. 5; Supplementary Tables S6-S9**; (Rasmussen et al., 2013; Balfagón et al., 2019). Interestingly, the *H. stipulacea* “Combination”-specific DEGs were also enriched in GO-terms related to defense against fungi, bacteria and insects (**Fig. 5; Supplementary Tables S6 and S9**). There are at least two possible explanations for the activation of defense-related genes. Firstly, abiotic and biotic stresses can affect subsequent plant responses both antagonistically and synergistically including the induction of biotic gene expression in response to abiotic stresses (Saijo and Loo, 2020). Secondly, seagrasses including *Halophila* sp. possess rich epiphytic and less characterized endophytic microbial communities (Garcias-Bonet et al., 2012; Conte et al., 2021; Tarquinio et al., 2021). Importantly, changes in environmental conditions can alter the community composition and presumably functional activity and may produce signals that are perceived by the seagrasses and induce expression of plant genes involved in plant-microbe interactions (Rotini et al., 2017; Szitenberg et al., 2022; Liu et al., 2023b).

### *H. stipilacea* plants from the anthropogenically-impacted TY site exhibit transcriptome responses to stress that could contribute to increased stress tolerance

Our data revealed significant differences in transcriptome responses between the seagrass populations from the pristine SB site and the anthropogenically-impacted TY site. Firstly, TY plants exhibited reduced transcriptome plasticity compared to SB plants across all stress treatments and time points (**Fig. 3; Supplementary Tables S2 and S3**). Plasticity refers to the ability of a single genotype to generate different phenotypes in response to varying environmental conditions. The extent of phenotypic change observed across these environments reflects the degree of plasticity (Kusmec et al., 2018; Schneider, 2022). Moreover this plastcity can be displayed across biologcal scales from phenotypic traits to gene expression. In terms of plant stress tolerance, plasticity is often compared between different genotypes or populations that have undergone selection for stress tolerance (Pigliucci et al., 1995; Swindell et al., 2007; Wang et al., 2020c; Pieri et al., 2024).While it is unknown whether the SB and TY populations represent different genotypes, a preliminary SNP analysis that we performed on nine *H. stipulacea* plants situated as little as 10 m from each other, revealed at least three genetically distinct groups (**Supplementary Fig. S3**). Since the SB and TY sites are approximately 3 km from each other, plants from these sites could represent genetically distinct populations.

An important question in evolutionary biology is whether there is any adaptive value to differences in phenotypic or gene expression plasticity (Rivera et al., 2021; Wang et al., 2022). Comparative transcriptome analysis of salt-mediated gene expression between *Arabidopsis thaliana* and its halophytic Brassicaceae relatives *Eutrema salsugineum* and *Schrenkiella parvula* has demonstrated adaptive value to differences in transcriptome plasticity that contribites to stress tolerance. The halophytes exhibit reduced transcriptome plasticity in response to ionic stress compared to stress-sensitive *Arabidopsis thaliana*, and this is partially related to the existence of excess NaCl- and boron-related “stress-ready” genes in *E. salsugineum* and *S. parvula*, respectively. In contrast, “stress-ready” genes are not found in *A. thaliana* (Kazachkova et al., 2018; Wang et al., 2021). “Stress-ready” genes involved in metal homestasis were also identified in the zinc hyperaccumlator and zinc-tolerant Brassicaceae species, *Arabidopsis halleri* (Becher et al., 2004). “Stress-ready” genes are characterized by high or low constitutive expression in extremophyte plants whereas the expression of orthologous genes in related stress-sensitive species is up/downregulated in response to stress. Similarly, we identified “stress-ready” genes in plants from the impacted TY site whereas such genes were not found in SB plants except for 2 genes in the “Combined” treatment (**Fig. 4; Supplementary Table S4**). Furthermore, these “stress-ready” genes were associated with reactive oxygen species and cell wall remodeling both of which are important in plant stress responses including in seagrasses (**Table 1; Supplementary Table S5**; Tenhaken, 2015; Tutar et al., 2017; Mittler et al., 2022; Pfeifer et al., 2022; Sandoval-Gil et al., 2023)). Notably, the largest proportion of TY “stress-ready” genes was found when we compared the response to the “Nutrient” treatment between SB and TY plants (**Fig. 4; Supplementary Table S4**). Over 90% of the genes assigned to the two response modes were “stress-ready” while less than 4% exhibited a “shared response” mode. A higher nutrient load is likely to be a primary difference between the pristine SB site and the anthropogenically-impacted TY site, and could act as a driver of changes at the molecular level (Pazzaglia et al., 2022). Although we only examined transcriptome profiles, the “stress-ready” state in the halophytic Brassicaceae is also reflected at the proteome and metabolome level (Kazachkova et al., 2018), and this might also be the case for TY plants. However, proteomic and metabolic profiling of the SB and TY populations in response to the three stressors would be necessary to confirm the “stress-ready” state of TY plants.

It should be noted that increased transcriptome plasticity in response to stress can also be linked to plant stress tolernace (Swindell et al., 2007; Eshel; et al., 2022). For instance in the Brassicaceae heat-tolerant desert species, *Anastatica hierochuntica* one third of the transcrtipome exhibits greater plasticity in response to heat stress than *A. thaliana* (Eshel et al., 2022). These highly heat-responsive genes are enriched in many biological processes related to abiotic stress. Thus, both reduced and elevated transcriptome plasticity can be associated with stress tolerance depending upon the species/genotype and envionmental context.

In addition to reduced transcriptome plasticity and the presence of “stress-ready” genes, the TY transcriptomes also exhibted greater recovery to near “Control” transcriptomes, 2 weeks post-stress, than SB transcriptomes (**Fig. 3b**). The capacity of plants to recover following stressful events often termed “stress resilience” is essential for resumption of growth and development after the stress event, and is being recognized as a critical trait in the study of stress tolerance (Li et al., 2020; Yuan et al., 2023; Bakery et al., 2024). At the molecular level, research using seagrasses was the first to link transcriptome resilience to stress tolerance (Franssen et al., 2011). In this seminal study, populations of *Zostera marina* from southern and northern Europe that exist in different thermal environments were subjected to a simulated heat wave that has led to actual mortality of the northern population. Transcritpome analysis revealed that gene expression patterns induced by stress were comparable in both populations. However, while the transcriptome in the southern population swiftly recovered to control values after the heat wave, the transcriptome of the northern population did not recover. Thus, the increased transcriptome resilience that we observed in the plants from the impacted TY site, might contribute to the stress tolerance of this population.

### *Core Stress-Response* (*CSR*) genes could act as molecular markers of early onset of stress in seagrass meadows

We further leveraged our transcriptome data to identify 38 *CSR* genes whose expression is up/downregulated by stress in both populations and in all stress treatments (**Fig. 6**). A putative role for the *CSR* genes in the core stress response machinery is supported by their functions in various abiotic and biotic stresses. Indeed, nine *CSR* genes are directly related to stress responses. In addition, two genes are involved in regulating the circadian clock, which controls the expression of many stress-responsive genes in an approximately 24-hour rhythm (Grundy et al., 2015). Expression of the *CSR* genes could act as early indicators of stress in the field before growth or mortality becomes evident and this gene set adds to a list of putative early stress indicator genes for *P. oceanica* (Santillán-Sarmiento et al., 2023). However, to employ *CSR* genes, it would be necessary to generate baseline stress-neutral expression levels in mesocosm experiments and/or in pristine seagrass meadows to assess whether the expression of a panel of these genes is above a certain threshold indicating that the seagrass meadow is under stress.

The importance of early stress indicator genes is emphasized by the startling findings from a seminal study by (Zandalinas et al., 2021), who challenged *Arabidopsis thaliana* with multifactorial stress combinations of six different abiotic stressors: salt, heat, high light intensity, oxidative stress, acidity and heavy metals. Each stress was applied so that, individually, they caused a minimal effect on plant growth. They reported that up to four stress combinations, there was a gradual deleterious effect on the plant growth. However, four or more stress combinations led to a drastic reduction in plant growth and survival. These data are alarming for seagrasses and marine organisms in general because of the increasing threat of multiple stressors impinging on aquatic ecosystems including eutrophication and associated effects (hypoxia, algae blooms, excess nutrient stress), changes in CO_2_ levels and acidification, extreme thermal events, and coastal chemical pollutants such as heavy metals, herbicides, and pharmaceuticals (Wilkinson et al., 2015; IOC-UNESCO, 2022; Li et al., 2023; Pinheiro et al., 2023; Glibert et al., 2024). Thus, the cumulative effect of these multifactorial stress combinations could have severe effects on seagrasses and their associated ecosystem services that are difficult to predict and challenging to detect.

In conclusion, we have shown that: (i) *H. stipulacea* plants from an anthropogenically-impacted site that are tolerant to single/combined thermal/excess nutrient stressors, possess a transcriptome with lower stress-induced plasticity, possesses “stress-ready” genes and which exhibits increased resilience compared to plants from the pristine SB site. These findings suggest that environmental conditions in seagrass habitats can drive local molecular adaptation; (ii) the response of seagrasses to combined stressors associated with climate change and coastal development cannot be predicted from the responses to single stressors; (iii) we have identified a panel of *CSR* genes that has the potential to serve as a useful early warning indicator of seagrass stress, which could be employed for management purposes, thereby preventing further decline of precious coastal aquatic ecosystems. In this respect, we emphasize that even a locally acclimated population might not be able to cope with ongoing ocean changes. Hence, we would strongly recommend stakeholders, policymakers and practitioners to better manage local stressors arising from climate change and coastal development in order to reduce the combined impact of local and global stressors on seagrasses.

## Acknowledgments

This work was funded by The Israeli Ministry of Science and Technology (MoST), Israeli-Italian Binational Grant Number 3-15152 (G Winters, PB-C), ICA in Israel, Grant 03-16-06a (G Winters), and the Goldinger Trust Jewish Fund for the Future (SB). This work was also financed by the SEANARIOS project (SEAgrass SCENARIOS under thermal and nutrient stress: FKZ 03F0826A), an Israeli-German Scientific Cooperation, funded by the German Federal Ministry of Education and Research (BMBF), jointly with the SEASTRESS project, an Israeli-Italian Scientific Cooperation, funded by the Ministry of Science and Technology of Israel (MOST). HMN was supported by Israel Science Foundation (ISF) grant no. 1015/21 (G Winters, SB) and a postdoctoral fellowship from the Jacob Blaustein Center for Scientific Cooperation. GP was supported by the European Union - NextGenerationEU (National Recovery and Resilience Plan (NRRP), Mission 4 Component 2 Investment 1.4, Project Code: CN00000033.We also would like to thank Tomás Azcárate-García for his support in experimental development and execution.

## Conflict of Interest Statement

The authors declare no conflicts of interest.

## Author Contributions

SB, G Winters and GP conceptualized and supervised the overall project; BY, PB-C and GW performed the mesocosm experiment. HMN and BY conducted the transcriptome analyses; MD and G Wang performed the SNP analysis. HMN and SB wrote the manuscript. G Winters, PB-C, and GP critically revised and approved the final manuscript.

## Data Availability

RNA-Seq reads are openly available via the NCBI SRA database under BioProject PRJNA1141374. Other main data that supports the findings of this study are available in the main text and Supplementary Information of this article. Commands and scripts used for data analysis are available upon request.

## Supplementary Information

**Supplementary Table S1:** Mesocosm nutrient levels for the duration of the experiment.

**Supplementary Table S2:** Raw read data plus Gene Ontology Terms and differentially expressed genes.

**Supplementary Table S3:** DEGs displayed in each category of Divenn diagrams (**Fig. 3**).

**Supplementary Table S4:** DEGs and transcript levels of “stress-ready” and “shared response” genes at 1 week of stress.

**Supplementary Table S5:** Gene Ontology Term enrichment of “stress-ready” and “shared response” genes at 1 week of stress.

**Supplementary Table S6:** Gene Ontology Term enrichment of combination-specific DEGs in SB plants after 1 week of stress, and gene lists for cluster 1.

**Supplementary Table S7:** Gene Ontology Term enrichment of combination-specific DEGs in TY plants after 1 week of stress, and gene lists for cluster 1.

**Supplementary Table S8:** Gene Ontology Term enrichment of combination-specific DEGs in SB plants after 5 weeks of stress, and gene lists for cluster 1.

**Supplementary Table S9:** Gene Ontology Term enrichment of combination-specific DEGs in TY plants after 5 weeks of stress and gene lists for cluster 1.

**Supplementary Figure S1:** Mesocosm temperature levels for the duration of the experiment.

**Supplementary Figure S2:** Global transcriptome response of genes associated with the GO-term “response to stress”, to single and combined thermal and excess nutrient stresses in SB (pristine site) and TY (impacted site) *H. stipulacea* plants.

**Supplementary Figure S3:** SNP analysis clusters individual *H. stipulacea* plants into three genetically distinct groups.

## Notes

### Competing Interest Statement

The authors have declared no competing interest.

### Summary of Updates

Current address of Dr. Hung Manh Nguyen was added.

