## Supplementary Figure S1 for "Transcriptome responses of two *Halophila stipulacea* seagrass populations from pristine and impacted habitats, to single and combined thermal and excess nutrient stressors, reveal local adaptive features and core stress-response genes" (from another version of this preprint: version 1, 2025-01-24)

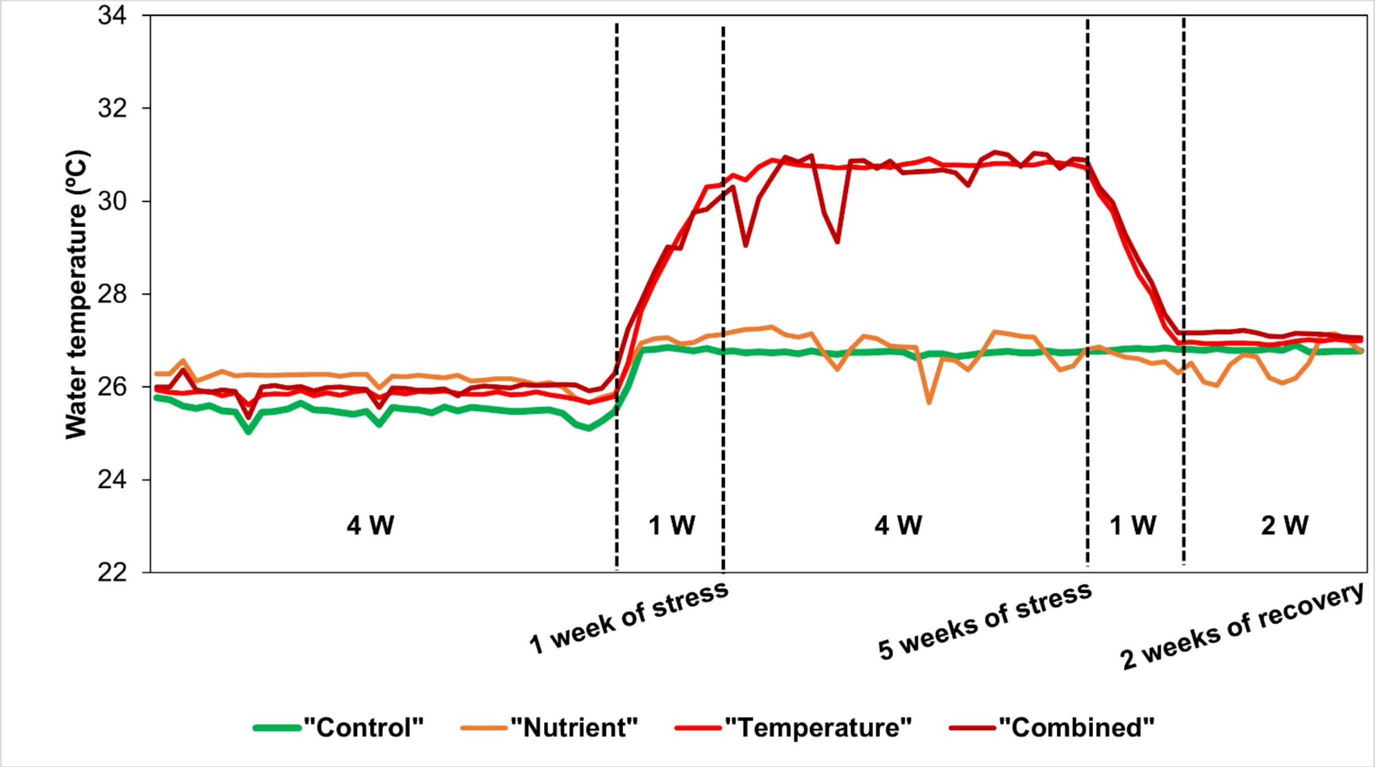


**Supplementary Figure S1.** Water temperature in aquaria during experimental period. Temperature readings were taken every hour and averaged to give the daily reading presented in the figure. Colored lines indicate aquaria assigned to control and each stress treatment. W, weeks.
