## Supplementary Figure S3 for "Transcriptome responses of two *Halophila stipulacea* seagrass populations from pristine and impacted habitats, to single and combined thermal and excess nutrient stressors, reveal local adaptive features and core stress-response genes" (from another version of this preprint: version 1, 2025-01-24)

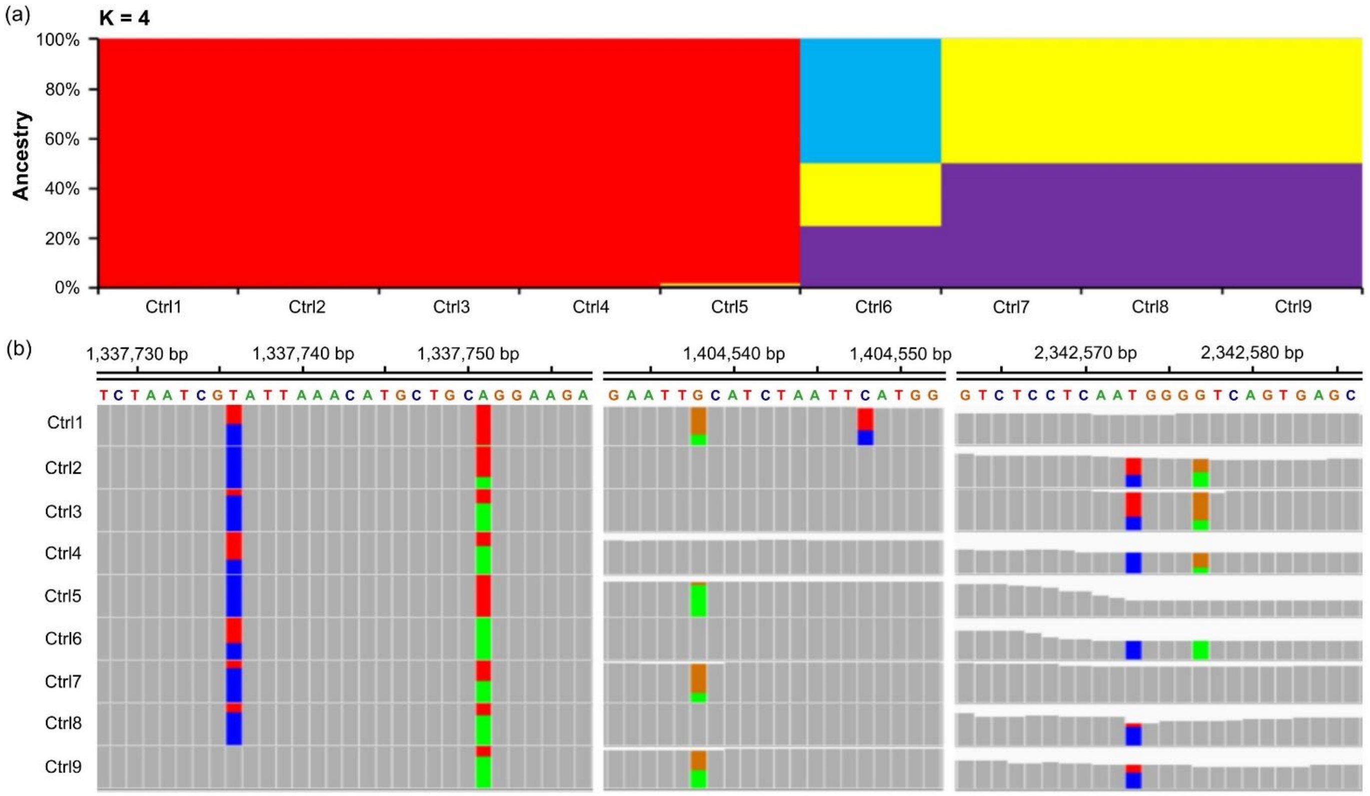


**Supplementary Figure S3.** SNP analysis clusters individual *H. stipulacea* plants into three genetically distinct groups. RNAseq reads were mapped to a 5 MB genomic region of the reference *H. stipulacea* using HISAT2 version 2.0.1 [1]. High confidence SNPs were identified using Samtools version 1.3 [2] with a predicted error rate less than 0.01 for a given SNP. We selected 679 biallelic SNP sites from a candidate pool of 695 SNPs to further examine the genetic diversity of the 9 samples using frappe version 1.1 [3] with population sizes (K) of 2-8 (shown is the population structure for K=4). Each SNP site was represented by at least 100 RNAseq reads uniquely mapped to the reference genome from all samples. (a) Clustering reveals three putative genotypes; (b) Examples of SNP alignment between samples. Each bar represents the read depths for a given site. Sites that show differences compared to the reference *H. stipulacea* genome are colored in each sample. Each color indicates a nucleotide, and bar height represents the proportion of reads with that variation.

1. **Kim D, Langmead B, Salzberg SL** (2015) HISAT: a fast spliced aligner with low memory requirements. Nat. Methods 12: 357-360.

2. **Li H, Handsaker B, Wysoker A, Fennell T, Ruan J, Homer N, Marth G, Abecasis G, Durbin R** (2009) The Sequence Alignment/Map format and SAMtools. Bioinformatics 25: 2078-2079.

3. **Tang H, Peng J, Risch NJ** (2005) Estimation of individual admixture: analytical and study design considerations. Genet Epidemiol 28: 289-301.
